# Defective DNA damage repair leads to frequent catastrophic genomic events in murine and human tumors

**DOI:** 10.1101/314518

**Authors:** Manasi Ratnaparkhe, John Wong, Pei-Chi Wei, Mario Hlevnjak, Thorsten Kolb, Daniel Haag, Yashna Paul, Frauke Devens, Paul Northcott, David TW Jones, Marcel Kool, Anna Jauch, Agata Pastorczak, Wojciech Mlynarski, Andrey Korshunov, Rajiv Kumar, Susanna M. Downing, Stefan M Pfister, Marc Zapatka, Peter J McKinnon, Frederick W Alt, Peter Lichter, Aurélie Ernst

**Author notes:** These authors contributed equally to this study (Manasi Ratnaparkhe, John Wong).

## Abstract

Chromothripsis and chromoanasynthesis are catastrophic events leading to clustered genomic rearrangements. Whole-genome sequencing revealed frequent chromothripsis or chromoanasynthesis (n= 16/26) in brain tumors developing in mice deficient for factors involved in homologous-recombination-repair or non-homologous-end-joining. Catastrophic events were tightly linked to *Myc/Mycn* amplification, with increased DNA damage and inefficient apoptotic response already observable at early postnatal stages. Inhibition of repair processes and comparison of the mouse tumors with human medulloblastomas (n=68) and glioblastomas (n=32) identified chromothripsis as associated with *MYC/MYCN* gains and with DNA repair deficiencies, pointing towards therapeutic opportunities to target DNA repair defects in tumors with complex genomic rearrangements.

---

Chromothripsis and chromoanasynthesis are two forms of genomic instability that lead to complex genomic rearrangements affecting one or very few chromosomes^1–3^. These two types of catastrophic events play a role in numerous tumor entities as well as in some congenital diseases^3,4^. The first form, chromothripsis, is characterized by the simultaneous occurrence of tens to hundreds of clustered DNA double-strand breaks^1,5^. The DNA fragments resulting from this shattering event are re-ligated by error-prone repair processes, with some of the fragments being lost. The outcome is a highly rearranged derivative chromosome, with oscillations between two or three copy-number states^6^. Conversely, the local rearrangements arising from chromoanasynthesis exhibit altered copy number due to serial microhomology-mediated template switching during DNA replication^2^. Resynthesis of fragments from one chromatid and frequent insertions of short sequences between the rearrangement junctions are associated with copy number gains and retention of heterozygosity^2^.

The availability of murine tumor models recapitulating these phenomena would substantially facilitate the investigation of the mechanistic aspects underlying chromothripsis and chromoanasynthesis. We showed previously the role of constitutive and somatic DNA repair defects in catastrophic genomic events in the context of *TP53* and *ATM* mutations^5,7^. Further factors essential to the biochemical and signaling context of occurrence of these catastrophic events remain to be identified, and the role of repair processes in chromothripsis and chromoanasynthesis needs to be better defined.

Homologous recombination repair (HR) and canonical non-homologous end-joining (cNHEJ) represent the two major repair processes for DNA double-strand breaks in mammalian cells. Conditional inactivation of key repair factors of either of these two pathways, such as BRCA2 (HR), XRCC4 or Lig4 (cNHEJ) in nestin-expressing or *Emx1*-expressing neural progenitor cells leads to medulloblastomas (MBs) or high-grade gliomas (HGGs) in mouse models with a p53-deficient background^8,9^ (McKinnon et al, in preparation). We now showed that these tumors frequently display chromothripsis/chromoanasynthesis – also in the absence of cNHEJ - and that analyzing tumor development in these mice will help us to understand the role of repair processes in catastrophic genomic events.

## RESULTS

### Inactivation of DNA repair factors essential for HR or cNHEJ leads to frequent chromothripsis and chromoanasynthesis

Whole-genome sequencing of the tumors developing in BRCA2/p53, XRCC4/p53 or Lig4/p53 deficient animals showed frequent complex genomic rearrangements (Figure 1a-c, Supplementary Figure 1, see Methods for details on the scoring criteria), with a prevalence of 64% in the XRCC4/p53-deficient mice (n=11 MBs), 60% in the Lig4/p53-deficient mice (n=5 HGGs), 71% in the BRCA2/p53-deficient HGGs (n=7), and 33% in the BRCA2/p53-deficient MBs (n=3) (Figure 1c). Most cases showed rearrangement patterns consistent with chromoanasynthesis, whereas two tumors in the BRCA2/p53-deficient animals were affected by chromothripsis. The high prevalence of catastrophic events in the tumors developing in these animals (see Discussion regarding the occurrence of such events when only p53 is inactivated) suggested that chromothripsis or chromoanasynthesis might represent driver events in tumorigenesis in these models. In line with this, murine orthologs of known MB and HGG oncogenes such as *Mycn*, *Myc*, *Cyclin D2* (in agreement with previous studies^8,9^) or *Braf* were frequently gained or amplified in association with the observed complex genomic rearrangements (Figure 1d, Supplementary Figure 1). These amplifications were linked with increased expression levels of the respective oncogenes as compared to the expression values observed in the normal brain (Supplementary Figure 2).

**Figure 1.**
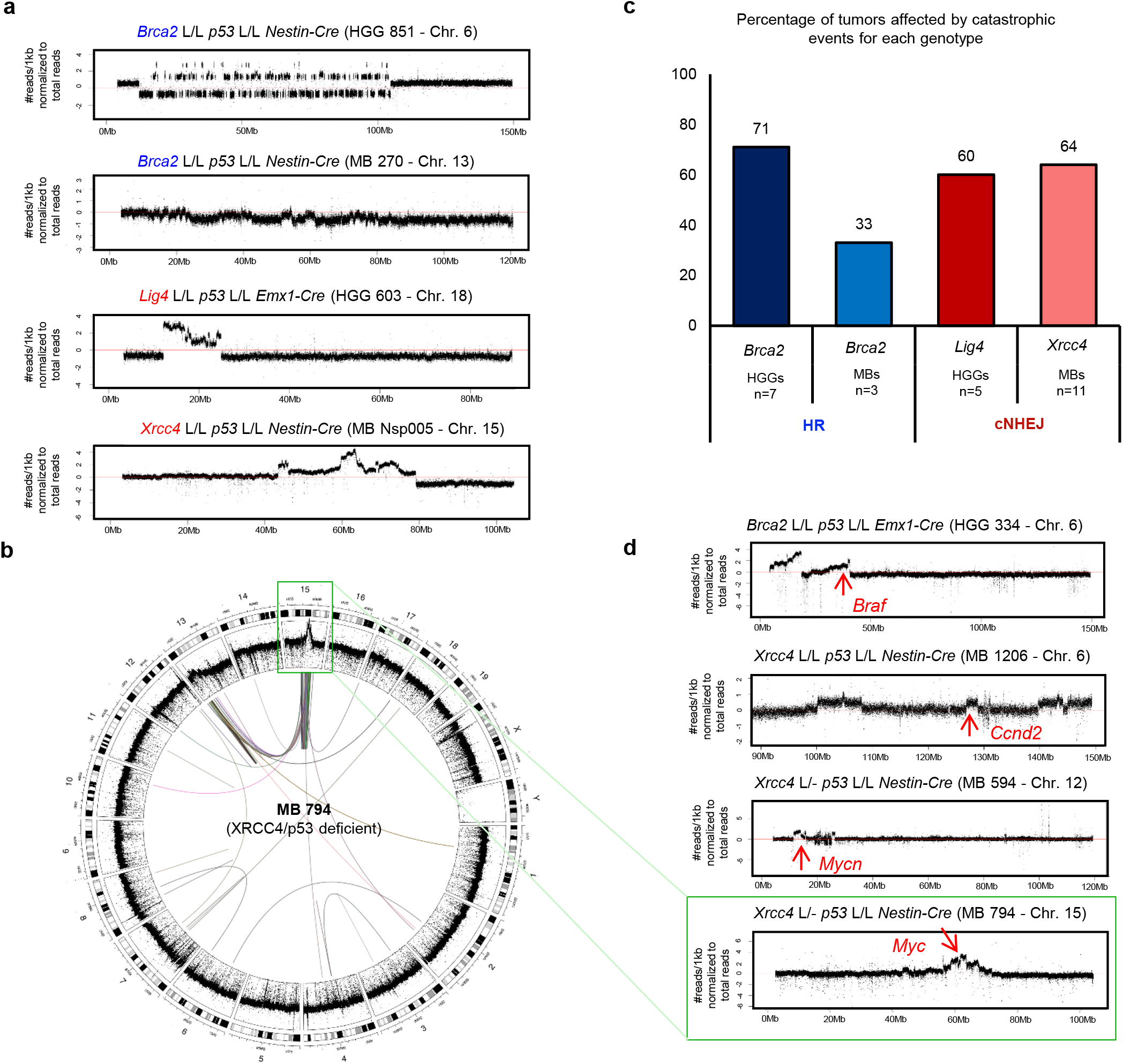
Inactivation of DNA repair factors essential for HR or cNHEJ in a p53 deficient background leads to frequent catastrophic events **a.** Examples of chromosomes affected by chromothripsis (upper plots) or chromoanasynthesis (lower plots) for one tumor for each mouse genotype. HGG, high-grade glioma; MB, medulloblastoma **b.** Circos plot for a medulloblastoma showing chromoanasynthesis on chromosome15 in a XRCC4/p53 deficient mouse. Magnified coverage plot for chromosome 15 is shown in d (lower panel) **c.** Prevalence for catastrophic events in brain tumors developing due to inactivation of HR (blue, Brca2/p53 deficient mice) or cNHEJ (red, Lig4/p53 or XRCC4/p53 deficient mice) in neural progenitors **d.** Chromothripsis and chromoanasynthesis drive tumor development, as shown by the amplification of oncogenes associated with the catastrophic events.

M-FISH analysis of MB cells from the XRCC4/p53 deficient mice showed that catastrophic events were associated with increased chromosome numbers (44 to 63 chromosomes per metaphase in tumors with chromoanasynthesis, as compared to 40 to 43 chromosomes per metaphase in tumors without chromoanasynthesis, Figure 2a). This finding is in line with the link between polyploidy and the detection of rearrangements due to one-off genomic catastrophic events^10,11^. In addition, tumor cells with chromoanasynthesis showed higher total numbers of aberrations and frequently harbored marker chromosomes (Figure 2a). Cases with several normal copies and one larger derivative chromosome (presumably the copy affected by chromoanasynthesis) suggest that polyploidization may precede chromoanasynthesis, facilitating the survival of the cell after such a catastrophic event. Interestingly, chromosomes affected by chromoanasynthesis were also shown to harbor recurrent breakpoints involved in translocations, deletions and amplifications as described previously^8^.

**Figure 2.**
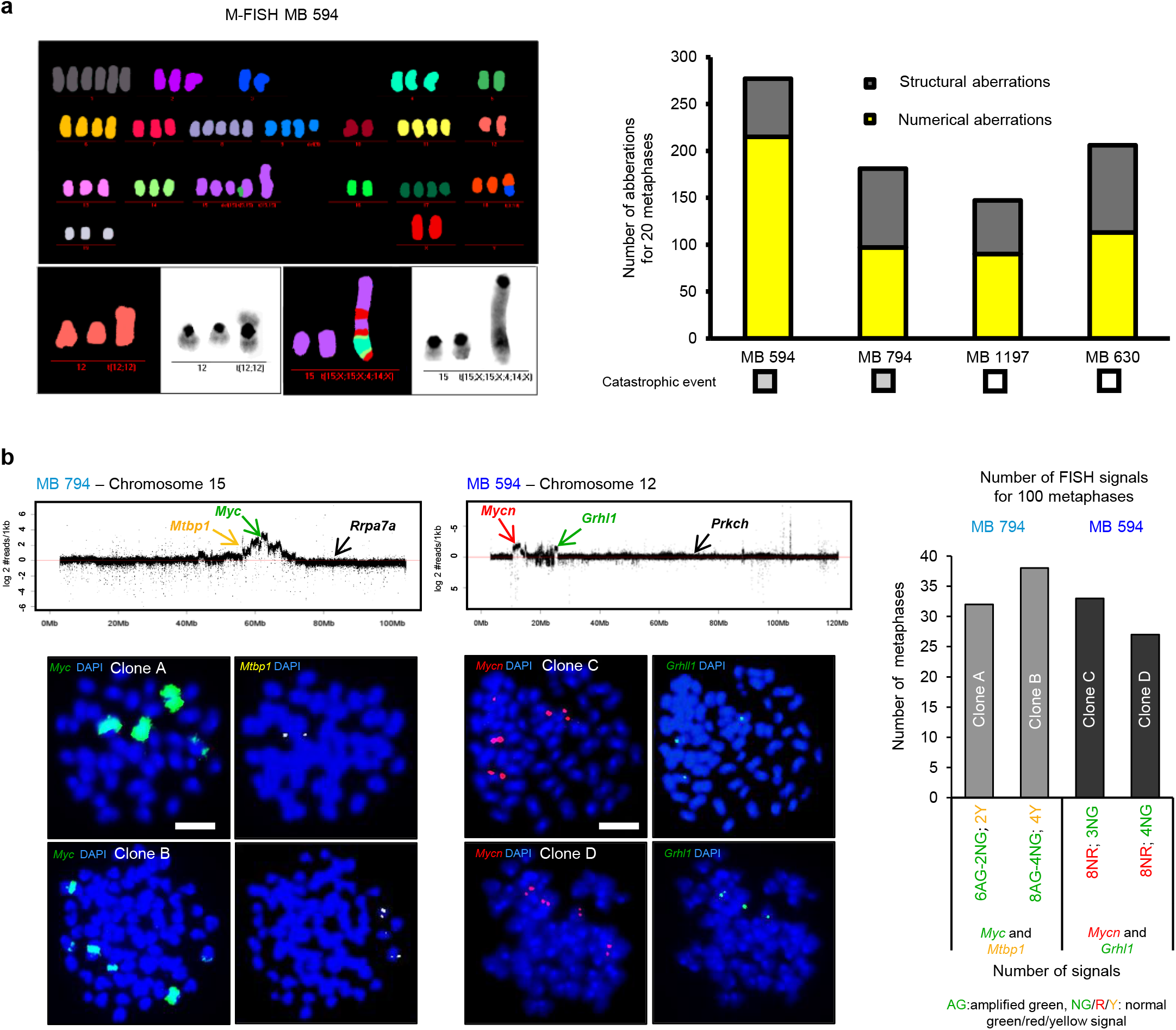
**a.** Chromoanasynthesis is associated with increased chromosome numbers and with structural aberrations. Left panel, M-FISH analysis of medulloblastoma cells from a XRCC4/p53 deficient mouse. Right panel, quantification of the structural and numerical aberrations detected by M-FISH analyses for medulloblastoma cells derived from four Xrcc4/p53 deficient mice **b.** Amplifications of *Myc* and *Mycn* likely facilitate catastrophic events. FISH analysis with probes for *Myc* (RP23 98D8, green) or *Mycn* (RP23 10C3, red) in combination with probes matching for loci affected by chromoanasynthesis (*Mtbp1*, RP23 288J22 and *Grhl1*, RP23 431C5) showed subpopulations of cells with amplifications of *Myc* or *Mycn* but without gain associated with chromoanasynthesis, suggesting that amplifications of *Myc* or *Mycn* occur before the catastrophic event. Quantifications of the signals for the control probes *Rpa7a* and *Prkch* are shown in Supplementary Figure 3. Magnification, X1000.

### Amplifications of *Myc* or *Mycn* likely facilitate catastrophic events in XRCC4/p53 mice

Strikingly, gains or focal amplifications of *Myc* or *Mycn* were detected in nearly all (10/11) MBs developing in the XRCC4/p53-deficient mice, in line with previous work (Table 1)^8^. From these, 6/11 independently showed both gains of *Myc* or *Mycn* and regions of chromoanasynthesis, whereby the *Myc* or *Mycn* loci were occasionally included in the region affected by the catastrophic event. In one tumor, we detected chromoanasynthesis but no gain of *Myc* or *Mycn*; however, the *Myc*n expression level in this MB was comparable to the level observed in the tumors with gains of *Mycn* (Supplementary Figure 2). There was no tumor with neither chromoanasynthesis nor gain of *Myc* or *Mycn*, suggesting that these two events play a driver role in tumor development in the context of cNHEJ and p53 deficiency in this entity. As gains of *Myc/Mycn* but no chromoanasynthesis were detected in 4/11 MBs, we hypothesized that gains of *Myc* or *Mycn* might be early events in the development of these tumors and that chromoanasynthesis possibly occurs later (in the remaining 6/11 cases), as one of the possible consequences of the activation of the oncogene (Table 1).

**Table 1.**

| MBs (in *Nestin-cre p53<sup>-/-</sup> Xrcc4<sup>-/-</sup>*) |  |  |  |
| --- | --- | --- | --- |
| MB ID | Chromo-anasynthesis | *Mycn* gain | *Myc* gain |
| 630 | No | yes | yes |
| 1187 | No | yes | no |
| 1197 | No | yes | no |
| 506 | No | yes | no |
| 1206* | Chr. 6 ,13 | no | no |
| 1207 | Chr. 7 | yes | no |
| 1224 | Chr. 15 | yes | yes |
| 706 | Chr. 7 | yes | no |
| 794 | Chr. 15 | yes | yes |
| 594 | Chr. 12 | yes | no |
| NXP005 | Chr. 13, 15 | no | yes |

\*Expression levels for *Mycn* and *Myc* were comparable to MBs where gains of these loci were detected

These observations prompted us to consider potential scenarios for the sequence of events between the gains of *Myc/Mycn* and chromoanasynthesis by two-colour FISH in two tumors (Figure 2b). In both tumors, two major clones were identified among the tumor cells, namely one clone with *Myc* or *Mycn* amplification, respectively, with co-occurrence of gains associated with chromoanasynthesis, and a second clone with *Myc* or *Mycn* amplification but no gain associated with chromoanasynthesis (Figure 2b, Supplementary Figure 3). The latter suggests that the amplification of *Myc* or *Mycn* presumably occurs before chromoanasynthesis and may facilitate this event, even though we cannot formally exclude the possibility of retention of the *Myc/Mycn* gain and a loss of the rearrangement due to chromoanasynthesis.

Gains of *Myc* or *Mycn* were not as frequent in the HGGs from the BRCA2/p53 (2/7) or Lig4/p53 (2/5) deficient animals (Supplementary Table 1), making an analysis of the sequence of events challenging. However, the correlation between catastrophic events and gains of *Myc* or *Mycn* holds true in these tumors as well, with all tumors showing a gain of either oncogene being affected by a catastrophic event.

### Increased DNA damage and inefficient apoptotic response may contribute to catastrophic events

In order to understand what leads to catastrophic events in the precursor cells, we analyzed the cerebellum from early postnatal stages up to tumor onset in age matched *Nestin-Cre p53* ^-/-^ *Xrcc4* ^c/c^, *Nestin-Cre p53* ^-/-^ *Xrcc4* ^c/+^ and *Nestin-Cre p53* ^-/-^ *Xrcc4* ^c/-^ animals (Figure 3). No difference was detected between the genotypes regarding the proportions of Ki67 positive cells in the cerebellum or regarding telomere features, as evaluated by immunofluorescence staining and by telomere FISH, respectively (Supplementary Figure 4). Loss of XRCC4 led to higher levels of DNA DSBs at all examined developmental stages, as assessed by gH2AX and 53BP1 staining (Figure 3, middle panels and Supplementary Figure 5). Therefore, alternative repair pathways seem to be unable to compensate entirely for the inactivation of cNHEJ in the neural tissue. Apoptosis was less frequent in the cerebellum of *Nestin-Cre p53* ^-/-^ *Xrcc4* ^c/-^ as compared to *Nestin-Cre p53* ^-/-^ *Xrcc4* ^c/+^ animals, as seen by TUNEL (Figure 3, lower panel and Supplementary Figure 5). The increased DNA damage together with the less efficient apoptotic response in neural tissue likely contribute to the frequent occurrence of catastrophic events in MBs developing in XRCC4/p53 deficient mice.

**Figure 3.**
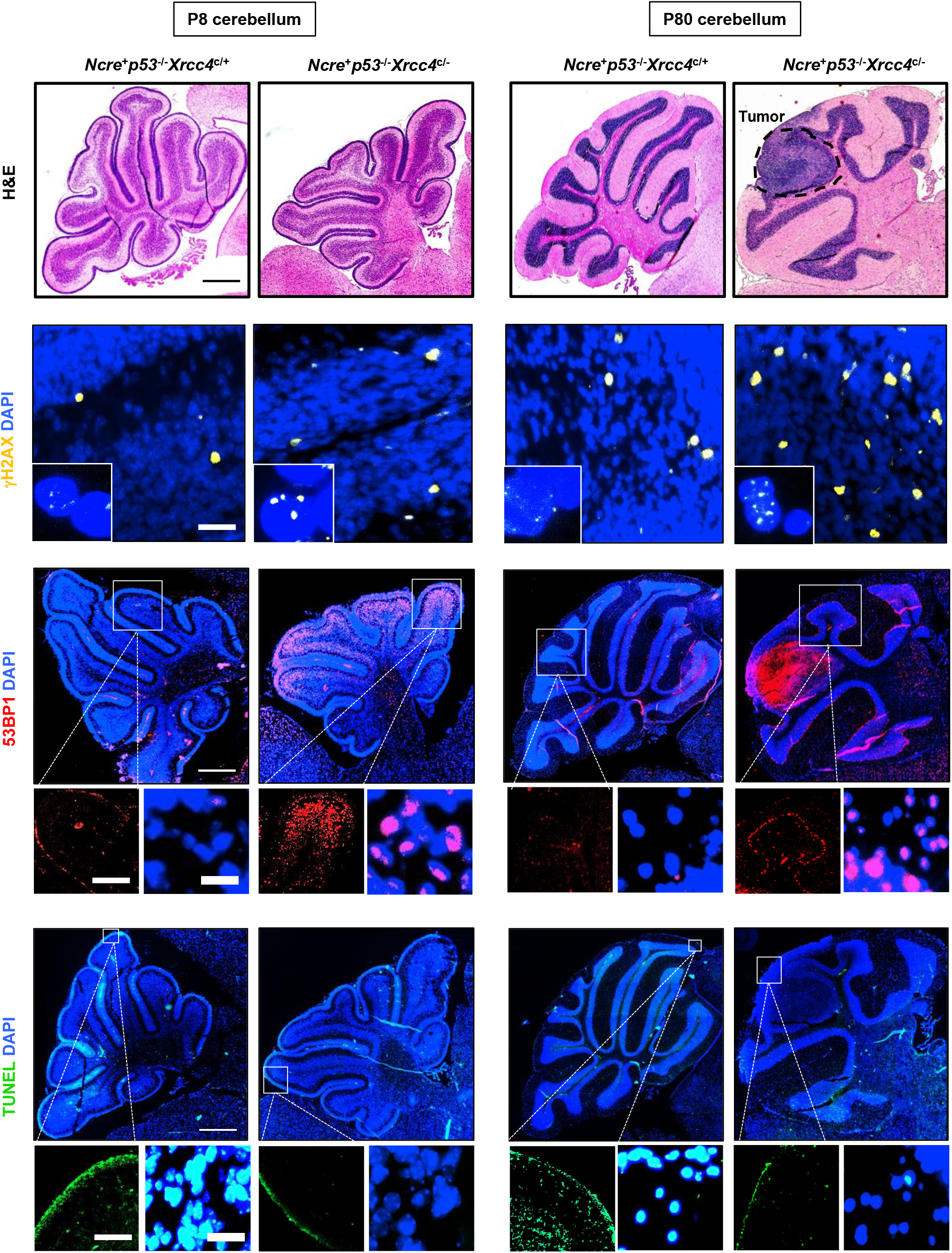
Analysis of brain tissue from early postnatal stages (P8) up to tumor formation (P80) in XRCC4/p53 deficient mice (*Nestin-cre p53* ^-/-^ *Xrcc4* ^c/-^) and control animals (*Nestin-cre p53* ^-/-^ *Xrcc4* ^c/+^). Upper panels, H&E staining of cerebellum sections shows the location of the tumor at P80 (magnification, X100). Middle and lower panels, representative immunofluorescence stainings for DSB markers gH2AX and 53BP1 and for terminal deoxynucleotidyl transferase (TdT)-mediated dUTP nick end labeling (magnification, X100; insets, X200 and X600). Representative images for 14analyzed cerebella are shown. Analysis of brain sections at intermediate stages (P60) are shown in Supplementary Figure 5.

Interestingly, loss of BRCA2 was reported to result in high numbers of DSBs in the cerebellum, as assessed by phosphorylated H2AX levels in *Brca2^LoxP/LoxP^;Nestin-Cre* and *Brca2 Nes-cre;p53^−/−^* animals, as compared to *Brca2^LoxP/LoxP^* control mice^12^. Apoptosis was described as widespread throughout the *Brca2^LoxP/LoxP^;Nestin-Cre* cerebellum, but attenuated when p53 is also inactivated^12^. Therefore, disruption of HR by deletion of *Brca2* in a p53 deficient background likely induces levels or types of DNA damage that cannot be repaired by other repair processes during neural development, similarly to what we observe in the context of cNHEJ deficiency.

### Repair processes involved in chromothripsis and chromoanasynthesis

In order to investigate the DNA repair processes involved in catastrophic events, we analyzed the expression levels of essential factors in the main repair pathways in the MBs developing in XRCC4/p53-deficient mice, in the non-neoplastic cerebellum and in granule neural progenitors (GNPs) of control animals of the same genotype (Figure 4a, left panel and Supplementary Figure 6). As XRCC4 is essential for efficient cNHEJ, this pathway is not functional in the *Xrcc4*-null neural progenitors, despite moderate levels of Lig4. This highlights the occurrence of catastrophic events in the absence of cNHEJ and raises the question of which other pathways may play a role in the repair processes.

**Figure 4.**
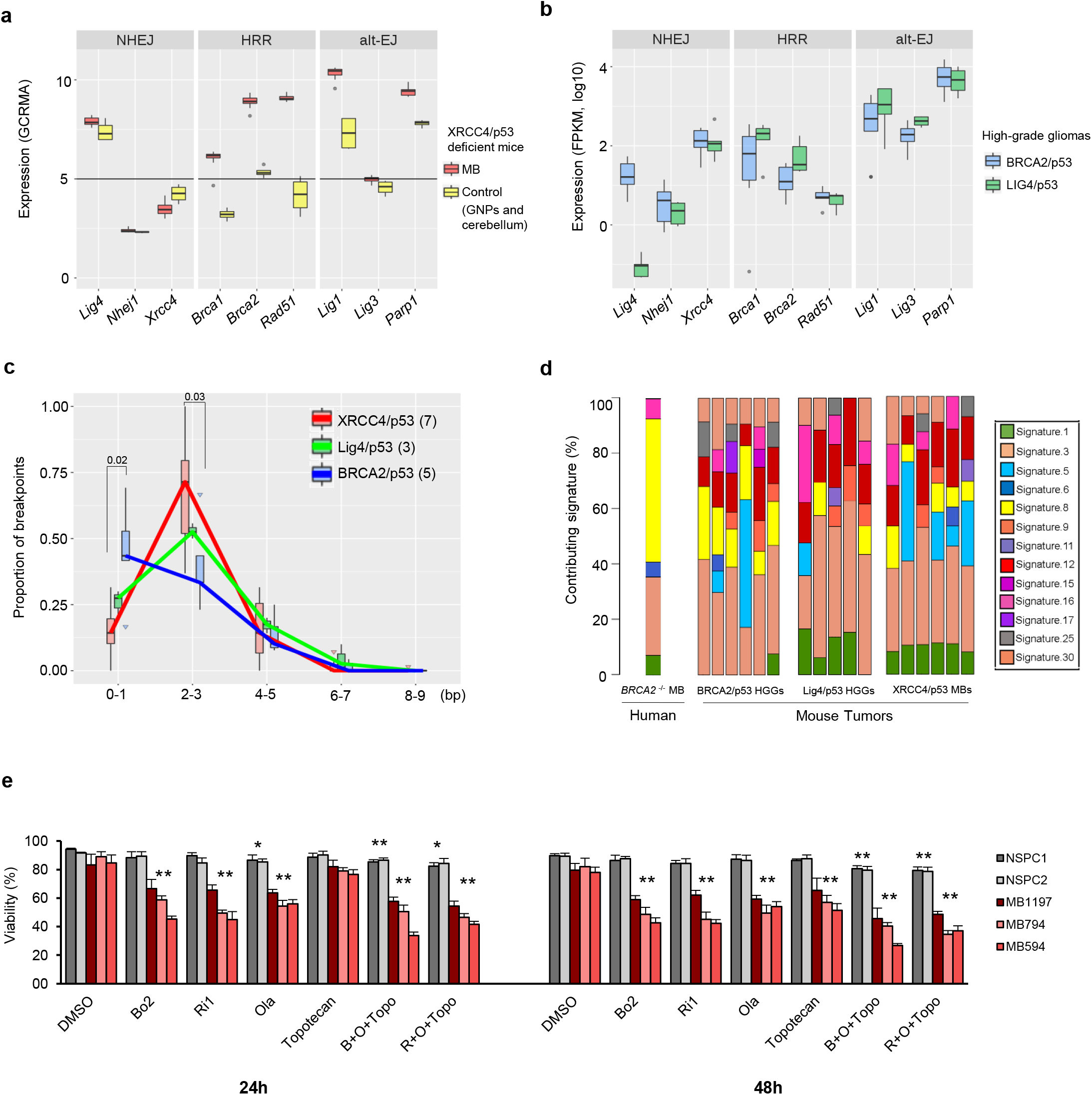
**a, b.** Normalized expression levels for repair factors involved in the main DNA DSB repair processes, namely cNHEJ, HRR and Alt-EJ. **a**. Red boxes show the expression levels in medulloblastoma cells of XRCC4/p53 mice (n=8) and yellow boxes show the expression levels in the healthy cerebellum and in granule neural progenitors (n=4) for age-matched animals of the same genotype (the expression levels for the non-neoplastic cerebellum and for granule neural progenitors were not significantly different and therefore shown as one box). The horizontal line depicts the threshold for which genes are considered to be expressed **b.** Normalized expression levels for repair factors in gliomas developing in BRCA2/p53 (blue boxes) or Lig4/p53 (green boxes) deficient animals **c.** Analysis of the length of the microhomologies at the breakpoints due to catastrophic events in the tumors developping in XRCC4/p53, Lig4/p53 and BRCA2/p53 animals. The analysis focuses on chromosome regions affected by the catastrophic events **d.** Mutational processes are dominated by signature 3 (associated with HR deficiency). Strikingly, the main contributing mutational signatures are very similar in tumors based on inactivation of factors essential for HR or for cNHEJ. A human MB from a *BRCA2* compound heterozygous patient is shown for comparison **e.** Synthetic lethality approaches targeting medulloblastoma cells from XRCC4/p53 deficient mice. Medulloblastoma cells (MB594, MB794 and MB1197) from *Nestin-Cre p53* ^-/-^ *Xrcc4* ^c/-^ and *Nestin-Cre p53* ^-/-^ *Xrcc4* ^c/c^ mice as well as control neural progenitor cells (NSPC1 and NSPC2) from *Nestin-Cre p53* ^-/-^ *Xrcc4* ^+/-^ mice were treated with RAD51 inhibitors to block HR (B02, 10uM or Ri1, 10uM), PARP inhibitor (Olaparib, 5uM), topotecan to induce DNA damage (100nM) or with combinations (B02 10uM, Olaparib 5uM and Topotecan 100nM or Ri1 10uM, Olaparib 5uM and Topotecan 100nM). Cell viability values are shown as averages +/- standard deviation for 3 independent experiments.

Key factors of HR were expressed at background levels in the normal cerebellum and in the GNPs but at moderate to high levels in the tumor cells (Figure 4a, middle panel). Alternative end-joining factors were expressed at moderate levels in the normal cerebellum and in the GNPs and at high levels in the tumor cells (Figure 4a, right panel), except Lig3, detected neither in the healthy cerebellum nor in the tumors in these animals. Consistent with this observation, Lig3 plays an essential role in mitochondrial DNA integrity but not in XRCC1-mediated nuclear DNA repair^13^. Altogether, as expected in the context of *Xrcc4* inactivation, HR at certain stages of the cell cycle (nearly absent in G1, most active in the S phase, and low in G2/M) but more likely alt-EJ are able to mediate the repair processes in the neural compartment of these animals. Interestingly, expression levels of repair factors in high-grade gliomas of BRCA2/p53 and Lig4/p53 deficient animals were very similar, with the exception of the two inactivated genes (Figure 4b).

As an additional approach to the expression analysis, detailed comparisons of the microhomologies at the breakpoint junctions on the chromosomes affected by chromothripsis or chromoanasynthesis allow the inference of which repair processes were presumably active at the time of the catastrophic event (Figure 4c, Supplementary Figure 6). In the tumors developing in the BRCA2/p53 deficient animals, we detected high proportions (close to 50%) of blunt ends and very short microhomologies (0 to 1 bp), consistent with a repair mediated by cNHEJ, which likely takes over in the absence of HR. For the XRCC4/p53 and Lig4/p53 animals, in contrast, the microhomologies were significantly longer (2 to 3 bp for 50 to 75% of the junctions), consistent with alternative end-joining and in agreement with previous studies^14^. This emphasizes the role of alternative end-joining in catastrophic events in the context of cNHEJ inactivation. Importantly, loss of p53 alone does not alter cNHEJ activity^14,15^.

Mutational signatures can be used to evaluate deficiencies in specific repair processes. By adapting the method developed by Alexandrov and colleagues to assign specific signatures to mutational processes in human tumors^16^, we identified mutational signatures contributing to somatic mutations in brain tumors developing in BRCA2/p53, Lig4/p53 and XRCC4/p53 deficient mice (Figure 4d). Interestingly, all tumors developing in a context of cNHEJ inactivation displayed contributing proportions of signature 3 (associated with a failure of DNA DSB repair by HR^17^) in the same range as tumors from BRCA2/p53 animals and from a human MB occurring in a patient with biallelic inactivation of *BRCA2*. The contribution of mutational signature 3 in the context of functional HR but inactive cNHEJ may point to a novel etiology for signature 3, namely consequences from excessive levels of DSBs and from the inability of the repair system to cope with high DNA damage, rather than necessarily pre-existing HR defect.

In order to show the dependence of the tumor cells on specific repair processes and to exploit DNA repair deficiencies, we treated the MB cells isolated from the *Nestin-Cre p53* ^-/-^*Xrcc4* ^-/-^ mouse tumors with inhibitors of HR (B02 or Ri1, RAD51 inhibitors) and/or Alt-EJ (olaparib, PARP inhibitor) and induced DNA damage using topotecan, a topoisomerase 1 inhibitor. As controls, we applied the same treatments to neural progenitor cells from age-matched heterozygous animals (*Nestin-Cre p53* ^-/-^ *Xrcc4* ^+/-^). Combinations of RAD51 inhibitor, PARP inhibitor and topoisomerase inhibitor dramatically decreased the viability of the MB cells without affecting the viability of the normal neural progenitor cells for short-term treatment (up to 48h, Figure 4e). For longer treatments, the viability of the neural progenitor cells was reduced at the tested concentration for the combination treatment, but the fraction of viable cells was still 30% higher than for the tumor cells (Supplementary Figure 7). Importantly, the doubling times of the control neural progenitor cells were not higher than those of the MB cells, showing that the observed difference in treatment response is not due to the respective proliferation rates.

### Catastrophic events in human brain tumors

To investigate the role of cNHEJ in catastrophic events in human neural cells, we used the CRISPR/Cas9 system to inactivate *TP53* and *LIG4* in iPSC derived neural progenitors. We induced DNA DSBs by topoisomerase inhibitor treatment and analyzed the levels of DNA damage and apoptosis by gH2AX staining and TUNEL, respectively. After CRISPR/Cas9-mediated disruption of *TP53* and *LIG4*, we detected similar levels of DSBs but considerably less apoptotic cells as compared to wild-type control cells and to p53-deficient/*Lig4* wild-type cells (Figure 5a and Supplementary Figure 8). Therefore, inactivation of DNA repair factors in combination with p53 deficiency might also contribute to catastrophic events in human neural precursor cells.

**Figure 5.**
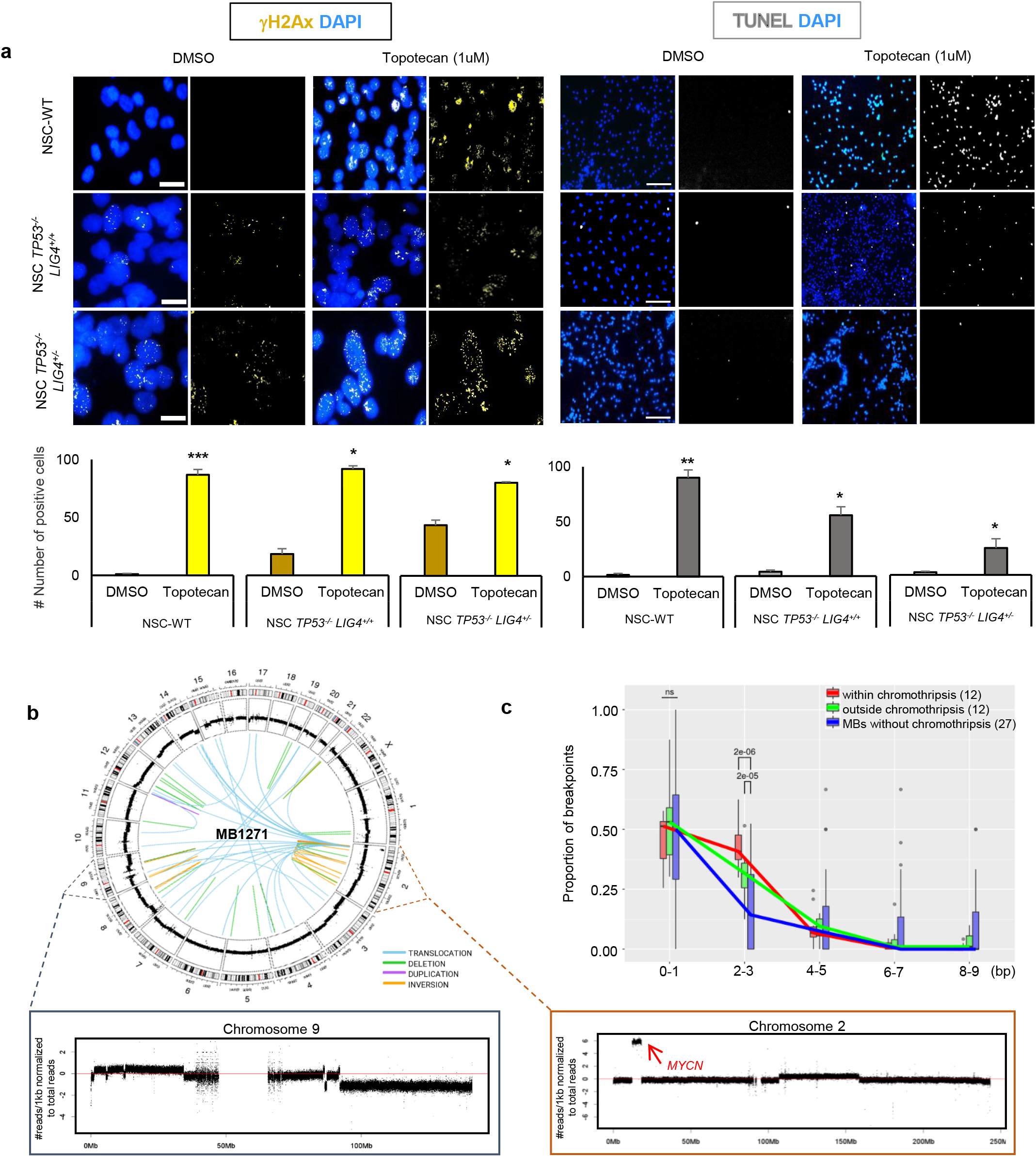
**a.** Response to DNA damage after CRISPR/Cas9 mediated disruption of *TP53* and *LIG4* in human iPSC-derived neural progenitor cells. Left panel, immunofluorescence analysis of DSB marker gH2AX. Right panel, TUNEL. Topotecan treatment (1uM) was applied for 48h to induce DNA DSBs. **b.** Chromothripsis in a human medulloblastoma with BRCA2 deficiency. Circos plot and coverage plots showing rearrangements associated with chromothripsis on chromosome 9 and *MYCN* amplification on chromosome 2. **c.** Analysis of the length of the microhomologies in human SHH MBs at the breakpoints due to catastrophic events (red), on chromosome regions not affected by catastrophic events in the same tumors (green) or genome-wide in MBs not showing chromothripsis (blue).

We next wanted to see how our findings on chromothripsis and chromoanasynthesis in the mouse tumors relate to catastrophic events detected in human primary brain tumors. We first sequenced a medulloblastoma from a patient with Fanconi anemia due to compound heterozygosity in *BRCA2* (c.657_658delTG and c.7558C>T). Interestingly, this tumor showed chromothripsis and focal amplification of *MYCN* (Figure 5b). Beyond rare cases with germline mutations, we re-analyzed whole-genome sequencing data from human SHH medulloblastomas^18^ (n=68) and pediatric glioblastomas^19^ (n=32). Consistent with the putative role of MYC and MYCN in catastrophic events that we identified in the murine XRCC4/p53 MBs, gains or focal amplifications of the *MYC* or *MYCN* loci were detected at higher frequencies in human tumors with chromothripsis as compared to cases without chromothripsis, both in glioblastomas and in SHH MBs (Supplementary Table 2). In line with this result, a link between *MYC* amplification and chromothripsis was previously shown in group 3 MB^20^. In the human MBs and GBMs as in the mouse tumors, the chromosomes affected by the catastrophic events were not necessarily the chromosomes harboring the *MYC* or *MYCN* locus, respectively (Supplementary Tables 1 and 2). Therefore, this association is not merely due to a selective advantage provided by the oncogene as a consequence of the massive rearrangement, but may in some instances be related to a facilitation of catastrophic events by *MYC* or *MYCN* activation. By a genome-wide search for amplified loci significantly associated with chromothripsis across 8 tumor entities, we identified the *MYC* locus as strongly associated with catastrophic events (Supplementary Figure 9).

Importantly, analysis of the microhomologies at the breakpoints showed high proportions of the longer microhomologies (2 to 3 bp for close to 50 % of the junctions) for breakpoints within the chromothriptic regions in human SHH MB (Figure 5c), consistent with alternative end-joining and in agreement with our findings on the murine tumors. In human pediatric glioblastomas, the fraction of junctions with longer microhomologies was also higher as compared to tumors without chromothriptic event, but did not reach the significance level (Supplementary Figure 10a). Altogether, these findings point towards a role for alternative end-joining in catastrophic events in human brain tumors.

At the mutational signature level, a significant association between chromothripsis and mutational signatures 3 and 8, associated with DNA repair defects, was reported in SHH medulloblastoma^11^. We confirmed the link between DNA repair deficiency and chromothripsis in additional tumor entities, namely adult glioblastoma (n=74), breast cancer (n=356) and melanoma (n=69) (Supplementary Figures 10 b-d). In glioblastoma, loss of *BRCA2*, *LIG4* or *XRCC4* was associated with more frequent occurrence of chromothripsis. In melanoma, we identified a significant association between mutational signature 3 (linked with HR deficiency) and chromothripsis. In breast cancer, high homologous recombination deficiency indexes were significantly associated with chromothripsis. Therefore, inactivation of HR and NHEJ factors is linked with chromothripsis across various tumor entities.

In analogy to the clinical use of PARP inhibitors in the context of BRCA-deficient breast cancer, our findings point towards therapeutic opportunities to target DNA repair defects in tumors with complex genomic rearrangements. Future studies will help defining the specific subsets of patients who may benefit from such a treatment.

## DISCUSSION

We showed that inactivation of factors of DNA DSB repair by HR or NHEJ leads to frequent catastrophic genomic events in murine and human tumors. The availability of murine models recapitulating recently discovered forms of genome instability will facilitate the investigation of the mechanistic aspects underlying one-off complex genomic rearrangements. It will be important to distinguish between the contribution of p53 and the role of the repair factors themselves. As *Nestin-Cre Xrcc4* ^c/-^ *p53* ^+/+^, *Nestin-Cre Xrcc4* ^c/-^ p53 ^+/-^, *Nestin-Cre Xrcc4* ^c/+^ *p53* ^+/-^*, Nestin-Cre Lig4 p53* ^+/+^ and *Nestin-Cre Lig4 p53* ^+/-^ animals do not die of medulloblastomas or gliomas^8,9^ it is hard to address this question in the same genetic background. The only study raising this issue showed rearrangements consistent with chromothripsis in MBs from *Ptch* ^+/−^; *p53* ^−/−^ mice and no chromothripsis in MBs from *Ptch* ^+/−^; *p53* ^+/+^ mice^5^. However, the tumors from the p53-deficient mice were all allografts from the same donor, and the group of *p53* wild-type mice had only three animals, therefore further work will be necessary. Even though deficiencies in repair factors in addition to the inactivation of p53 likely enhance the probability for catastrophic events, evidence from direct comparisons will be essential.

In humans, chromothripsis was detected across a wide range of tumor entities, whereas chromoanasynthesis was mainly described in the context of congenital diseases^2,4^. Here, we detected predominantly rearrangements resembling the outcomes of chromoanasynthesis and rarely events consistent with the definition of chromothripsis (see Methods for details on the scoring procedures). This discrepancy in the prevalence of both types of catastrophic events between human and mouse tumors may relate to inter-species differences in terms of development of the neural lineages in the context of DNA repair deficiency, number of cell divisions, extent and type of exposure to external and internal triggers (e.g. replication stress), telomere biology, sensitivity to DNA damage or efficiency of specific repair processes and apoptotic response. Another potential explanation for this divergence may relate to differences in scoring methods and definitions for catastrophic events, and especially the fact that chromoanasynthesis was described later as compared to chromothripsis, implying that some of the initially reported chromothriptic events might now be referred to as chromoanasynthesis.

Interestingly, we observed extreme examples of catastrophic events with exceptionally high numbers of breakpoints (see Figure 1a) and other cases with lower numbers of breakpoints. Further work will be needed to understand whether the former are extreme outcomes of the same process, which are lethal for the cell in the vast majority of cases. Alternatively, the degree of the catastrophic event may also relate to the affected cell type, as the nestin compartment comprises a large population of cells in the developing brain. Depending on the specific cell affected by the event, the brain region, the time point, the consequences may be tolerated or not, giving an advantage to a cell in a particular context.

We showed in the mouse models that amplifications of *Myc* and *Mycn* may facilitate catastrophic events, and not represent a consequence of these in all cases. Driving cell proliferation in the context of repair deficiency likely enhances the risk of chromosome fragmentation (e.g. as a consequence of chromosome segregation errors during mitosis and DNA damage on the missegregated chromosome) as well as DNA replication errors and collapsed replication forks, both processes being proposed to explain the rearrangements in chromothripsis and chromoanasynthesis^21^. The role of MYC and MYCN in replication stress^22^ may represent a causative factor in the occurrence of catastrophic events. Interestingly, *MYCN* is a frequent driver gene in SHH MBs, which represent the molecular subgroup of MBs developed by Li-Fraumeni Syndrome patients, for which all analyzed tumors show chromothripsis^5^. Both for HGGs and for SHH MBs, the frequency of cases with chromothripsis displaying gains of the *MYC* or *MYCN* loci was higher than in the tumors where no chromothripsis was detected, pointing to a role of this oncogene in catastrophic events.

Surprisingly, we observed very similar mutational signatures between tumors developing in BRCA2/p53, Lig4/p53 and XRCC4/p53 deficient animals. In particular, mutational signature 3, described as associated with defects in HR, was detected in tumors from Lig4/p53 and XRCC4/p53 mice. Even though we cannot exclude a previously unrecognized role for XRCC4 and Lig4 in HR, the more likely explanation is that signature 3 may be linked with different types of repair defects, including deficiency in cNHEJ, or associated with an exposure to extreme levels of DNA damage exceeding the DSB repair capacities. This mutational signature may point to mutational processes taking place when the major DSB repair pathways are saturated, not necessarily inactive due to pre-existing HR defects. If it is validated in human tumors, this result may have important clinical implications, given the relevance of HR repair defects for therapeutic targeting, in particular for the use of PARP inhibitors based on BRCAness.

Altogether, our findings on the tight links between DNA repair deficiencies and catastrophic events and on synthetic lethality approaches bear the potential to identify targets for new therapeutic strategies for tumors with complex genomic rearrangements.

## ONLINE METHODS

### Library preparation and sequencing

Purified DNA was quantified using the Qubit Broad Range double-stranded DNA assay (Life Technologies, Carlsbad, CA, USA). Genomic DNA was sheared using an S2 Ultrasonicator (Covaris, Woburn, MA, USA). Whole-genome sequencing and library preparations were performed according to the manufacturer’s instructions (Illumina, San Diego, CA, USA or NEBNext, NEB). The quality of the libraries was assessed using a Bioanalyzer (Agilent, Stockport, UK). Sequencing was performed using the Illumina X Ten platform. Information on available sequencing data for all cases are summarized in Supplementary Table 1.

### Cell Culture

NSPC1 and NSPC2 cell lines were grown as neurospheres and cultured in growth medium NBBG -Neurobasal A (Thermo Fisher, 10888-022), 2% B27- RA (Thermo Fisher, 12587-010), 0.5 mM Glutamax (Thermo Fisher, 35050061), 5 ug/mL Gentamicin (VWR International, SAFSG1397) with 10 ng/mL EGF (Thermo Fisher, PHG0315), FGF (Thermo Fisher, PMG0035) and PDGF (Thermo Fisher, PMG0045). *Nestin-Cre p53*-/- *Xrcc4*-/- medulloblastoma cells derived from the murine tumors for MB 594, MB 630, MB 794, MB 1197, MB 1206, MB 1207, MB 1224 and NxP005 were all cultured and maintained as previously described^23^. All related animal work was performed under protocol 14-10-2790R approved by the Institutional Animal Care and Use Committee of Boston Children’s Hospital. iPSC derived neural progenitors were cultured and maintained as previously described^24^.

### M-FISH

M-FISH for cell lines MB 594, MB 630, MB 794 and MB 1197 was performed as described previously^25^. Briefly, seven pools of flow-sorted human chromosome painting probes were amplified and directly labeled using seven different fluorochromes (DEAC, FITC, Cy3, Cy3.5, Cy5, Cy5.5 and Cy7) using degenerative oligonucleotide primed PCR (DOP-PCR). Metaphase chromosomes immobilized on glass slides were denatured in 70% formamide/2xSSC pH 7.0 at 72°C for 2 minutes followed by dehydration in a degraded ethanol series. Hybridization mixture containing labeled painting probes, an excess of unlabeled cot1 DNA, 50% formamide, 2xSSC, and 15% dextran sulfate were denatured for 7 minutes at 75°C, pre-annealed at 37°C for 20 minutes and hybridized at 37°C to the denaturated metaphase preparations. After 48 hours the slides were washed in 2xSSC at room temperature for 3x 5 minutes followed by two washes in 0.2xSSC/0.2% Tween-20 at 56°C for 7 minutes, each. Metaphase spreads were counterstained with 4.6- diamidino-2-phenylindole (DAPI) and covered with antifade solution. Metaphase spreads were recorded using a DM RXA epifluorescence microscope (Leica Microsystems, Bensheim, Germany) equipped with a Sensys CCD camera (Photometrics, Tucson, AZ). Camera and microscope were used with the Leica Q-FISH software and images were processed on the basis of the Leica MCK software and presented as multicolor karyograms (Leica Microsystems Imaging solutions, Cambridge, United Kingdom).

### NICK Translation and Two-color FISH

Nick translation was carried out for BAC clones all obtained from Source Bioscience of Myc (RP23 98D8), MycN (RP23 10C3), Grh1l (RP23 431C5) and Mtbp1 (RP23288J22). Two-color FISH^26^ was performed on metaphase spreads of MB 594 and MB 794 using fluorescein isothiocyanate-labeled probes and rhodamine-labeled probes. Pre-treatment of slides, hybridization, post-hybridization processing and signal detection were performed as described previously. Samples showing sufficient FISH efficiency (>90% nuclei with signals) were evaluated. Signals were scored in, at least, 100 non-overlapping metaphases. Metaphase FISH for verifying clone-mapping position was performed using peripheral blood cell cultures of healthy donors as outlined previously.

### H&E staining, immunofluorescence and TUNEL on tissue sections

Hematoxylin and eosin (H&E) staining and immunofluorescence stainings were performed on 4 μm formalin-fixed paraffin-embedded mouse brain sections. H&E staining was evaluated by a neuropathologist. Tissue sections from each cohort of p8, p60 and p80 for at least 3 animals per genotype were stained. Sections were deparaffinized, antigen retrieval was performed in 10mM citrate buffer pH 6.0 for 40 minutes and sections were cooled down to room temperature. Slides were then washed with PBS for 5 min and then blocked with blocking solution (10% goat serum diluted in PBS, 0.2% Triton-X) for 1 hour. Slides were then incubated with primary antibodies, γh2Ax (Cell Signaling, 9718S) 53BP1 (Santa Cruz, sc-22760), Ki67 (Abcam, 15580) overnight at 4°C. The next day slides were washed 3 times with PBS for 10min each and incubated for an hour at room temperature with rabbit secondary (Invitrogen, Alexa 488, A11034 or Life Technology, Alexa 568, A11036) at 1:100 dilution. After the incubation, slides were washed with PBS three times for 10 min each and then were mounted with DAPI Fluromount (Southern Biotech, 0100-020) and image analysis of the mouse cerebellum was done with Axio Zeiss Imager.M2 microscope. For TUNEL, the manufacturer’s protocol for paraffin embedded slides was followed (Roche, 11684795910).

### Telomere FISH

Telomere FISH (Agilent, K5325) was performed according to the manufacturer’s protocol as well. Slides incubated with the probe were washed 3 times 10 min each with PBS and then were incubated with secondary antibody Alexa 568 (Thermo Fischer, A11036) at 1:1000 dilution for 1 hour at room temperature. Slides were then washed again with PBS for 10 min each 3 times and were mounted with DAPI Fluoromount (Southern Biotech, 0100-020). Image analysis was done with Axio Zeiss Imager.M2 microscope.

### Synthetic Lethality

Tumor cell lines MB 594, MB 794, MB 1197 and control cell lines NSPC1 and NSCP2 were plated in 6-well plate for 10 hours and then treated either with 10uM B02 (Sigma Aldrich, SML0364) or 10uM Ri1 (Merck Millipore, 553514) or 5uM Olaparib (Biozol, AZD2281) or 100nM Topotecan (Apex Bio, B4982) or a combination of B02, Olaparib and Topotecan or a combination of Ri1, Olaparib and Topotecan at the above mentioned concentrations for 24 hours, 48 hours or 72 hours. For each time point, cell viability was measured by cell viability analyzer Vi-Cell XR (Beckman Couter).

### CRISPR-Cas targeted gene disruption

Guide RNAs for *TP53* and *LIG4* were constructed and cloned into lenti CRISPR v-2 (Addgene, 52961) according to the original online protocol of the Zhang lab (http://www.genome-engineering.org/crispr/wp-content/uploads/2014/05/CRISPR-Reagent-Description-Rev20140509.pdf). Following genes were targeted: *TP53* (gRNA: GT GAG GCT CCC CTT TCT TG), *LIG4* (gRNA: GAC GGA AAA GAT ACC TCG G). Virus production and transduction as described previously^27^; in brief, pLentiV2, pDMDG.2, and pSPAX were co-transfected in HEK293T cells, and virus-containing supernatant was concentrated by ultra-centrifugation. Transduction was done by adding concentrated virus particles to both NSC cell lines for 24hours, after which cells were maintained under selection either with puromycin or blasticidin at 2ug/mL for at least 2 weeks. After selection, cell lysates were made for western blotting or cells were grown for further experiments.

### Western Blotting

Western blot experiments for Alpha-Tubulin (Abcam, 52866), LIG4 (NovousBioNBP2-16182), TP53 (Progen, 65139) was carried out on NSC cell line protein lysates as described^7^.

### Immunofluorescence for cultured cells

For immunofluorescence, as previously described^7^, cells were grown on coverslips in 6-well plate and treated with Topotecan (1uM) or DMSO (control) for 48 hours, fixed and stained for γH2AX, 53BP1 or TUNEL (according to manufacturer’s protocol) and imaged with Axio Zeiss Imager.M2 microscope. For each of the three independent experiments 100 cells were counted per staining to score for positive cells.

### Bioinformatic analysis

#### Processing of short reads data

All whole genome sequencing data were aligned and processed by the DKFZ OTP pipeline.

#### Detection of somatic structural variants

Svaba was used to detect the breakpoints of the structural variants. The tool utilizes the sequence assembly approach for the detection of structural variants and InDels. For microhomology analysis, breakpoints with precisely mapped junctions were extracted from the output. Different approaches were taken for the germline subtraction to retrieve somatic structural variants depending on the availability of germline specimens, either A) matched germline control or B) pooled controls were used. The pooled control strategy is only used for highly inbred mice samples. Human samples without reliable germline controls were excluded from the analysis.

#### Microhomology analysis for structural variants

Svaba provides the microhomology information for each precisely mapped rearrangement. The analysis was focused on blunt end junctions or junctions with microhomologies 1-9bp in length. Junctions fulfilling these criteria were extracted and further grouped into 5 different categories (blunt-1bp, 2-3bp, etc). Structural variants within the chromothriptic region and outside the chromothriptic region were compared to assess whether there are significant differences in their usage of homology sizes.

Extracted breakpoint sets with less than 5 breakpoints fulfilling all abovementioned criteria in were excluded from the analysis.

#### Somatic InDels analysis

Somatic InDels were identified by Svaba. Blacklists for highly discordant repetitive regions for mouse genome mm10 were created by the joint calling of 25 mouse matched normal specimens in the mice cohort. The blacklist was used to filter out InDels that are likely to be false positives in the tumor-normal paired detection mode in Svaba. The sequence context of the small deletions was checked by samtools to evaluate if they carry microhomologies, small insertions were excluded from the analysis. For each sample, InDels were divided into two sets, which separate InDels within chromothriptic regions from InDels outside chromothriptic regions. Two sets were then compared by grouping into 9 different size bins (1-9bp) to evaluate the differences in each of their abundance.

#### Expression array analysis

The R library `affy` was used to process the raw CEL files of the Affymetrix Mouse 430 v2.0 expression array. To facilitate expression level comparison between genes, GCRMA was used for the normalization of probe-set expression values. GCRMA considers the sequence information of different probes to adjust for the binding affinity bias. Intensity values from multiple probes on the same gene were combined by averaging. GCRMA value of 5 was considered as the baseline of gene expression in the analysis.

#### Mutation calling for the identification of mouse and human mutational signature

Somatic nucleotide variations were identified in whole genome sequencing data using Mutect2, part of the GATK 4.0.1.2 bundle. Mutect2 involves two Bayesian classifiers to identify high confidence candidate somatic mutations at a given genomic locus using tumor and matched normal BAM files. Resulting VCF files with non-db SNP variants is used as input for calling cancer signatures in mouse.

Supervised mutational signature analysis of high-confidence somatic SNVs in individual samples was performed using non-negative matrix factorization formalism as described previously^11,16^. The expanded set of 30 canonical mutational signatures was used for decomposition of somatic mutations (http://cancer.sanger.ac.uk/cosmic/signatures). Furthermore, canonical mutational signatures were re-normalized using the observed trinucleotide frequency in the mouse genome to the one of the human genome for mouse samples.

## Data availability

The datasets generated and/or analyzed during the current study are available from the corresponding author on reasonable request.

## Acknowledgements

We would like to thank Michaela Hergt and the tissue bank of the National Center for Tumor Diseases (NCT, Heidelberg, Germany) for technical assistance, Achim Stephan for help with library preparation, Laura Siebert and Norman Mack for advice on paraffin embedding, Andrea Wittmann for help with the preparation of metaphase spreads, and Brigitte Schoell for M-FISH analyses. We would also like to express our gratitude to Angela Schulz, Nicolle Diessl, Laura-Jane Behl, Stephan Wolf from the DKFZ Genomics & Proteomics Core Facility, to Katja Beck and to the Data management group for excellent support with the next-generation sequencing analyses. Marc Zuckermann and Emma Phillips are gratefully acknowledged for advice in the planning of the CRISPR experiments, Dominik Sturm for help with re-analysis of TCGA data and Susanne Gröbner for kindly helping with mutational signature analyses. We also thank Marius Wernig for kindly sharing the iPSC line 6-a (06C53141). We would like to express our special appreciation and thanks to Pierre-Olivier Frappart, Natalia Voronina, Kendra Maaß, Daisuke Kawauchi and Scott Pomeroy for discussions, and to Claus Bartram and Magnus von Knebel-Doeberitz for advice. Finally, we would like to thank Michael Hain for excellent IT support, and the Fritz Thyssen Stiftung for financial support.

## Author contributions

MR performed most of the experiments; JW, MH and YP performed bioinformatic analyses; PCW performed the animal experiments with the Xrcc4/p53 mice; TK contributed to the CRISPR/Cas experiments; DH derived neural progenitors from human iPSCs; FD contributed to the FISH and sequencing experiments; RK provided the melanoma sequencing data; AP and WM contributed to the collection of human samples; AK performed neuropathology evaluation of tumor specimen; AJ analyzed the M-FISH data; DTWJ, MK PN contributed to data analysis; SMD provided RNA-seq data; SMP, MZ, PJMK, FWA, PL and AE contributed to the original concept of the project and experimental design.

---

**Supplementary Figure 1**

Coverage plots for all analyzed murine tumors

**Supplementary Figure 2**

Chromothripsis and chromoanasynthesis drive tumor development

**Supplementary Figure 3**

Controls for two-colour FISH for *Myc* and *Mycn*

**Supplementary Figure 4**

Telomere FISH and immunofluorescence analysis for Ki67 in the brain of Xrcc4/p53 deficient and control heterozygous mice

**Supplementary Figure 5**

TUNEL and 53BP1 staining at intermediate developmental stages

**Supplementary Figure 6**

Expression levels for main repair factors and controls for microhomology analysis (indels)

**Supplementary Figure 7**

Synthetic lethality in mouse cells: late time point (72h)

**Supplementary Figure 8**

Validation on CRIPSR Cas9 experiment, Immunofluorescence for 53BP1

**Supplementary Figure 9**

Pan cancer analysis of the association between chromothripsis and *MYC* status in human tumors

**Supplementary Figure 10**

Microhomology at the breakpoints in human GBM and Pan cancer analysis of the association between repair deficiencies and chromothripsis

**Supplementary Table 1**

Summary of the mouse tumors analyzed in this study, including scoring for catastrophic events and status of *Myc* and *Mycn*

**Supplementary Table 2**

Link between chromothripsis and gains of *MYC* or *MYCN* in human SHH MB and GBM

